# Vaccination with a Replication-Dead Murine Gammaherpesvirus Lacking Viral Pathogenesis Genes Inhibits WT Virus Infection

**DOI:** 10.1101/2024.11.20.624603

**Authors:** Dipanwita Mitra, Darby Oldenburg, J. Craig Forrest, Laurie T. Krug

**Affiliations:** HIV and AIDS Malignancy Branch, National Cancer Institute, Bethesda, MD, USA; Gundersen Medical Foundation: Virology Research, La Crosse, WI, USA; Department of Microbiology and Immunology, University of Arkansas for Medical Sciences, Little Rock, Arkansas, USA

**Keywords:** gammaherpesvirus, MHV68, vaccine, replication-dead virus, latency establishment, reactivation

## Abstract

Gammaherpesviruses are oncogenic pathogens that establish lifelong infections. There are no FDA-approved vaccines against Epstein-Barr virus or Kaposi sarcoma herpesvirus. Murine gammaherpesvirus-68 (MHV68) infection of mice provides a system for investigating of gammaherpesvirus pathogenesis and testing vaccine strategies. Prime-boost vaccination with a replication-dead virus (RDV) that does not express the essential replication and transactivator protein (RTA) encoded by *ORF50 (*RDV-50.stop) protected against WT virus replication and reduce latency in C57BL/6 mice and prevented lethal disease in *Ifnar1-/-* mice. To further improve the RDV vaccine and more closely model KSHV vaccine design, we generated an RDV lacking the unique M1-M4 genes and the non-coding tRNA-miRNA-encoded RNAs (TMERs) 6, 7, and 8 that collectively promote latency of MHV68 *in vivo*. Prime-boost vaccination of mice with RDV-50.stopΔM1-M4 elicited neutralizing antibodies and virus-specific CD8 T-cell responses in lungs and spleens, the respective sites of acute replication and latency, that were comparable to RDV-50.stop vaccination. When challenged with WT MHV68, vaccinated mice exhibited a near-complete block of lytic replication and a reduction in latency and reactivation. We conclude that major determinants of MHV68 pathogenesis are not required components for eliciting a protective immune response.

## Introduction

Kaposi sarcoma herpesvirus (KSHV; human gammaherpesvirus 8) and Epstein–Barr virus (EBV; human gammaherpesvirus 4) are cancer-causing gammaherpesviruses (GHV) that establish chronic persistent infections. KSHV is the causative agent of multiple cancers including the angioproliferative Kaposi sarcoma (KS) that manifests at cutaneous and visceral sites and two B cell lymphoproliferative disorders, primary effusion lymphoma and a variant of multicentric Castleman disease. The seroprevalence of KSHV is >40% in Sub-Saharan Africa (SSA), 10-30% in Mediterranean regions and <10% in northern Europe, Asia and the USA [1]. KSHV infection is a significant health burden in people living with HIV (PLWH) world-wide and in endemic areas including SSA [2]. KSHV associated cancers are a leading cause of morbidity and mortality in PLWH even when HIV is well controlled. Salivary oral transmission occurs in endemic regions, whereas the route of transmission in non-endemic areas, particularly among gay, bisexual, and other men who have sex with men, is less clear [3]. Blood transfusions and tissues transplants from KSHV-positive donors can also lead to infection [4,5]. EBV is ubiquitous and infects more than 90-95% of the adult population worldwide. EBV infections can cause infectious mononucleosis (IM), B-cell lymphomas [6], epithelial-cell cancers [7], and is etiologically associated with the development of multiple sclerosis [8,9]. EBV is primarily transmitted through saliva, but it can also spread via breast milk, other bodily fluids, and transplantation of EBV-positive organs [6].

GHV infection in the host is comprised of lytic replication and periods of quiescent latency. Lytic replication may occur upon initial infection at mucosal sites, followed by a latent program of limited viral gene expression that evades host immune detection [10,11]. Reactivation from latency to the lytic phase occurs sporadically and leads to the production of infectious viral progeny, allowing the virus to infect other cellular reservoirs within the host and to spread to new hosts [11]. Once latency is established, infection is incurable, and individuals are at lifelong risk of cancer from GHV infections [1]. Immune suppression increases the incidence of GHV cancers and indicates that immune control is key to prevention [12].

There is no FDA-approved vaccine against GHVs. Nucleoside analogs reduce lytic infection, but these do not impact the chronic, latent phase infection. A prophylactic ferritin nanoparticle conjugated glycoprotein 350 (gp350) subunit vaccine (NCT04645147) [13] and an mRNA vaccine containing gp350, gH/gL, and gp42 (NCT05164094) [14] are in clinical trials for EBV [15]. There is no KSHV vaccine in clinical trials. An effective vaccine strategy is needed to prevent infection and virus-associated cancers in populations most at risk for disease. One of the major challenges for KSHV vaccine design is the lack of knowledge regarding immune correlates of protection [16]. KSHV-specific T cell responses are weaker compared to EBV in infected individuals [17]. Additionally, there is a paucity of animal models available to investigate these systems *in vivo*. Efforts are underway to address this gap in the field, including the recent development of a humanized mouse model of KSHV/EBV co-infection [18], but this model does not recapitulate all aspects of an antigen-driven immune response and does not support virus infection of non-lymphoid cells.

Murine gammaherpesvirus 68 (MHV68; murid herpesvirus 4) infection of mice is a model pathogen system with direct genetic and biological correlates to KSHV that is applied to study GHV pathogenesis and define immune correlates of protection [10,19]. For both KSHV and MHV68, lytic replication and reactivation is controlled by the replication and transcription activator protein, RTA, encoded by open reading frame 50 (*ORF50*) [20,21]. Insertion of a premature translation stop codon in *ORF50* results in a replication-dead virus (RDV-50.stop) that infects cells but is unable to replicate [21]. We produced a high-titer, revertant-free virus stock in an *ORF50* codon shuffled-producer cell line [22]. Inoculation with RDV-50.stop exposes the host to intact virion particles and enables limited viral gene expression upon *de novo* infection, without production of infectious particles [23]. Prime-boost intraperitoneal (IP) vaccination of C57BL/6 mice with RDV-50.stop elicited neutralizing antibodies and virus-specific effector T cell responses in the lungs and spleen, respective sites of acute replication and latency. When challenged intranasally with WT MHV68, mice vaccinated with RDV-50.stop exhibited a near-complete block in lytic viral replication in lungs at 7 days (d7) post-challenge, and a significant reduction in latency establishment and reactivation from latency in spleen at d16 post-challenge. RDV-50.stop vaccination also protected immunodeficient *Ifnar1^−/−^* mice from lethal disease upon WT MHV68 challenge [23].

To more closely model KSHV vaccine design and further attenuate the MHV68 RDV-50.stop vaccine platform, the removal of genes unique to MHV68 genes should be considered. A previous study characterized a spontaneous 9.5 kb-deletion mutant of MHV68 (GHV68Δ9473) lacking the M1, M2, M3, and M4 genes in addition to eight small hybrid non-coding RNAs that encode tRNA like-element linked to miRNAs, termed TMERs. This spontaneous mutant exhibits a reduction in latency establishment and reactivation from latency in splenocytes [24]. Results from these studies corroborate with another murine gammaherpesvirus isolate, MHV76, with a slightly larger 9,538 bp deletion that encompasses M1-M4 and the 8 TMERs. Upon intranasal infection of mice, MHV76 exhibited a defect in acute replication and increased infiltration of inflammatory cells in the lungs, as well as decreased splenomegaly and establishment of splenic latency. Despite this defect at early phases of infection, MHV76 established long-term latency in both lungs and spleen [25].

The left-end coding and non-coding genes of MHV68 broadly promote pathogenesis *in vivo*. MHV68 M1 encodes a superantigen-like factor that stimulate Vβ4+ CD8 T cells to suppress virus reactivation via IFNγ production [26]. M2 encodes an adapter protein that facilitates latency establishment and reactivation from latency in B cells [27,28], augments B cell survival and proliferation [27], and promotes MHV68 dissemination to distal latency reservoirs [29]. M2 is a target for CD4 and CD8 T cells with cytotoxic functions during latency [30,31]. M3 encodes for a chemokine-binding protein that inhibits antiviral and inflammatory responses during acute and latent phases of infection [32–35]. M4 encodes a secreted glycoprotein that promotes splenic latency establishment [36]. The TMERs broadly promote B cell latency in immunocompetent mice [37,38] and virulence in immunocompromised mice [39].

Given that the genetic elements in the M1-M4 locus play critical pathogenic roles in immune modulation, latency and reactivation, we sought to test if loss of these genes would impact the protective immune response generated by an RDV vaccine candidate. Toward this end, we generated a second generation RDV candidate that lacks nucleotides 2,022 to 9,739 to remove M1, M2, M3, M4 and TMERs 6, 7 and 8. Here, we test the efficacy of this second generation RDV vaccine to generate an immune response and protect against WT MHV68 infection in mice.

## Materials and Methods

### Cells and Viruses

Primary C57BL/6 murine embryonic fibroblasts (MEFs) and NIH 3T12 (ATCC CCL-164) were cultured in Dulbecco’s Modified Eagle Medium (DMEM) supplemented with 10% fetal bovine serum (FBS), 100 IU/ml penicillin, 100 ug/ml streptomycin, and 2 mM L-glutamine to form complete DMEM (cDMEM). Cells were maintained in a standard humidified incubator at 37°C in 5% CO_2_. Primary MEFs at passages 2 and 3 were used for limiting dilution reactivation assays.

Viruses utilized in this study include WT MHV68 (WUMS strain) for vaccine challenge experiments and comparative immune responses in C57BL/6 mice and bacterial artificial chromosome (BAC)-derived [40] replication-deficient RDV-50.stop and RDV-50.stopΔM1-M4 recombinant MHV68 whose recombineering are described below. RDV-50.stop and RDV-50.stopΔM1-M4 recombinant MHV68 were produced on codon-shuffled RTA (CS-RTA)-expressing NIH 3T12 cells [22] and were validated by plaque assay on vector control cell lines to demonstrate absence of WT revertants.

Oligonucleotide sequences used for recombineering and sequencing are described in Table S1. To mutate the ORF50 locus of MHV68, two gBLOCKS (IDT DNA, Coralville, IA) were designed such that there was 30 bp overlap on the 5’ end of the first gBLOCK (50_STOP_FS_gBLK_1) and the 3’ end of the second gBLOCK (50_STOP_FS_ gBLK_2). Similarly, to delete the M1-M4 region of MHV68, two gBLOCKS, M1_4DEL gBLK_1 and M1_4DEL gBLK_2 were designed with a 30 bp overlap. This overlap was engineered to facilitate Gibson assembly using the HiFi Assembly mix (New England Biolabs, Ipswich, MA). The assembled gBLOCKs were diluted 1:4 and 2 uL were used for a PCR using the following primers to generate the full-length targeting construct: 50_STOP_FS_FOR and 50_STOP_FS_REV for RDV-50.stop; M1-4_DEL_FOR and M1-4_DEL_REV for RDV-50.stopΔM1-M4.

The 50_STOP or M1-M4_DEL PCR-amplified targeting construct was gel-purified and electroporated into the *En passant* E. coli strain GS1783 [41] harboring either WT MHV68 BAC or the ORF50.stop mutant BAC clone, respectively. Kanamycin -resistant colonies were chosen for resolution of the kanamycin-resistance cassette that is facilitated by I-SceI digestion and red recombination. Kanamycin-sensitive clones were analyzed for mutation of the ORF50 locus by PCR and subsequent Sanger sequencing of the PCR product, using the primers: 50_FSS_SEQ_FOR and 50_FSS_SEQ_REV. Kanamycin-sensitive clones were analyzed for loss of the M1-M4 region by PCR using the following primers: M1-4_SEQ_FOR and M1-4_SEQ_REV. Clones yielding a 1318 bp PCR product indicated that the M1-M4 region was successfully deleted. Clone #121 (RDV-50.stop) and clone #466 (RDV-50.stopΔM1-M4) were subject to full genome sequencing via Illumina based sequencing (Seq Center, Pittsburgh, PA). The resulting reads were analyzed using the Galaxy server and compared to the wild type MHV68 sequence (Genbank: U97553.2) to confirm that the intended M1-M4 region was deleted and the ORF50 frameshift/stop mutations were present. No other differences in the viral genome sequences were observed, excluding repetitive elements.

### Animal Studies

Female C57BL/6 mice were purchased from Charles River Laboratories (Wilmington, MA). Eight to ten-week-old mice were anesthetized using 1-4% isoflurane in an induction chamber and intraperitoneally (IP) inoculated with 1 X 10^6^ PFU of RDV-50.stop or RDV-50.stopΔM1-M4 diluted in 500 μL cDMEM and boosted with an equivalent dose on d28 post-prime. Sham-vaccinated mice inoculated with cDMEM only and RDV vaccinated C57BL/6 mice were challenged by intranasal (IN) infection with 1 × 10^3^ PFU of WT MHV68 diluted in 20 μL cDMEM on d28 post-boost. At the indicated timepoints, blood, lung, and spleen tissues were harvested following humane euthanasia using isoflurane or CO2.

### Pathogenesis assays

For acute replication titers, mice were euthanized with isoflurane at 7 days post-infection (dpi), and both lungs were removed and frozen at −80 °C prior to disruption in 1 ml of 10% cDMEM using 1 mm zirconia beads in a bead beater (Biospec, Bartlesville, OK). NIH 3T12 cells were plated at a density of 1.8 x 10^5^ cells/mL 1 day prior to infection. Serial dilutions of lung homogenate were overlaid on the NIH 3T12 cells for 1 hr at 37 °C, with rocking every 15 min, followed by an overlay of 5% methylcellulose (Sigma, St. Louis, MO) in cDMEM and incubated at 37 °C. After 7–8 days, cells were fixed with 100% methanol (Sigma) and stained with 0.1% crystal violet (Sigma) in 20% methanol, and plaques were scored.

For limiting dilution analysis, spleens were homogenized in a TenBroek tissue disrupter. Red blood cells (RBC) were lysed by incubating in 8.3 g/L ammonium chloride for 5 min at room temperature with inversion and then diluted with 25 mL cDMEM prior to centrifugation. Cells were filtered through a 70- or 100-micron filter prior to enumeration and downstream analyses. Limiting-dilution (LD)-PCR analyses to quantify frequencies of latently infected splenocytes were performed as previously described [42]. Briefly, 3-fold serial dilutions of latently infected cells were diluted in a background of uninfected NIH 3T12 fibroblasts. After overnight digestion with proteinase K, cells were subjected to a nested PCR targeting the ORF50 region of the viral genome. Single-copy sensitivity and the absence of false-positive amplicons were confirmed using control standards. Amplicons were visualized by SYBR Safe DNA gel stain (Thermo Fisher Scientific, Waltham, MA) in 1.5% agarose gels upon electrophoresis.

*Ex vivo* reactivation efficiency was determined as previously described [43]. Briefly, 2-fold serial dilutions of latently infected splenocytes were plated on MEF monolayers for analysis of reactivation from splenocytes from C57BL/6 mice. The presence of preformed infectious virus was detected by plating mechanically disrupted cells on indicator monolayers in parallel. Cytopathic effect was scored d14 and d21 after plating on MEFs.

### Flow cytometry

Spleens were processed using a tissue grinder and RBC lysis as described above. Lungs were digested using 150 U/ml Collagenase Type IV (Millipore Sigma, Burlington, MA) and 10 U/ml DNaseI for 2 hrs at 37°C with intermittent vortexing, followed by RBC lysis. 2 X 10^6^ single cell suspensions prepared from the lung or spleen were resuspended in 200 μl of PBS with 2% fetal bovine serum and blocked with TruStain fcX (BioLegend, San Diego, CA). T cell subsets were identified with antibodies against CD45 (dilution 1:200; clone 30-F11; BV510; cat# 563891) purchased from BD Biosciences (Franklin Lakes, NJ), TCRβ (dilution 1:200; clone H57-597; PerCP-Cy5.5; cat# 109228), CD8 (dilution 1:400; clone 53-6.7; BV785; cat# 100750), CD62L (dilution 1:200; clone MEL-14; FITC; cat# 104406), CD127 (dilution 1:200; clone 678 A7R34; PE-Dazzle594; cat# 135032), and KLRG1 (dilution 1:200; clone 2F1/KLRG1; PE-Cy7; cat# 138416) purchased from BioLegend, San Diego, CA and CD44 (dilution 1:200; clone IM7; APC-eFluor780; cat# 47-0441-82) purchased from eBioscence (Thermo Fisher Scientific). 1:5 dilution of H-2K(b)-p79 (cat# JD02150) or H-2D(b)-p56 (cat# JA02153) MHC-peptide dextramer complexes conjugated to PE or APC purchased from Immundex (Fairfax, VA) were added directly to antibody-stained samples. The data was collected on a CytoFLEX flow cytometer (Beckman Coulter, Brea, CA) and analyzed using FlowJoX v10.10.0 (Treestar Inc., Ashland, OR). CD8+ T cells were first gated as singlets, then live per exclusion of Alexa Fluor™ 700 NHS Ester, followed by gating for CD45+ and lymphocytes based on forward and side scatter parameters, and then as CD8+ TCRβ.

### Peptide stimulation

For analysis of effector T cell responses, 1 x 10^6^ splenocytes were plated into each of five wells of a 96-well flat-bottom plate and either left untreated or treated with 1 ug/ml of p56 (AGPHNDMEI) or p79 (TSINFVKI) peptides (Genscript, Piscataway, NJ) for 5 h at 37 °C in the presence of Brefeldin A (BD Cytofix/Cytoperm, BD Biosciences). The Fc receptors were blocked prior to surface staining with antibodies against CD45, TCRβ CD8, CD4 (dilution 1:600; clone GK1.5, BV711; cat# 100447; BioLegend), CD4 (dilution 1:200; clone GK1.5; AF488; cat# 100423; BioLegend) and CD44. Upon fixation and permeabilization with the BD Cytofix/Cytoperm kit (BD Biosciences), cells were stained with antibodies to IFNγ (dilution 1:100; clone XMG1.2, PE-Cy7, cat# 505826) and TNFα (dilution 1:100; clone MP6-XT22, BV650, cat# 506333, BioLegend). CD8+ T cells were first gated as lymphocytes based on forward and side scatter parameters, singlets, then live per exclusion of Alexa Fluor™ 700 NHS Ester, followed by gating for CD8+CD45+, and then CD44^hi^.

### Neutralization assay

Serum was prepared by collecting blood from sham-vaccinated or vaccinated mice, followed by centrifugation at 21,100 x g for 20 mins, after which the separated serum was carefully collected. Neutralization was tested by a plaque reduction neutralization test (PRNT) [23]. Briefly, three-fold serum dilutions, starting from an initial concentration of 1:80 in cDMEM, were incubated with 50 PFU of MHV68 at 37 °C for 1 hr. The virus/serum mixture was then added to a sub-confluent NIH 3T12 monolayer (1x 10^5^ cells/well) plated the previous day in a 12-well plate, in duplicate. As a control, 2 wells received no-serum added virus. Infected cells were overlaid with methylcellulose and incubated at 37 °C for 6-7 days. Methylcellulose media was then aspirated, and cell monolayers were fixed and stained as described above. Percent neutralization was determined by comparison of the number of plaques in experimental wells compared to no-serum added control wells, and each data point was the average of three wells.

### Statistical analysis

All data was analyzed using GraphPad Prism software (Version 10, La Jolla, CA). Frequencies of immune cells were analyzed by one-way ANOVA followed by post-tests depending on parametric distribution, as indicated. Total number of an immune cell subset per animal was log_10_-transformed prior to ANOVA. Titer data were log_10_-transformed and then analyzed with unpaired t-test or one-way ANOVA for multiple groups. Based on the Poisson distribution, the frequencies of viral genome–positive cells and reactivation were obtained from the nonlinear regression fit of the data where the regression line intersected 63.2%. The log_10_-transformed frequencies of genome-positive cells and reactivation were analyzed by unpaired, two-tailed t-test or one-way ANOVA for multiple groups.

## Results

### Vaccination with RDV-50.stopΔM1-M4 induces virus-specific adaptive immunity in mice

RDV-50.stop and RDV-50.stopΔM1-M4 mutants were constructed using a BAC recombineering approach [40]. Our previous study used an ORF50.stop mutant virus with a translation stop codon at amino acid 116 in the RTA protein [21,23]. To facilitate recombineering of additional mutations we recreated the RDV-50.stop mutant in the MHV68 BAC by introducing a frameshift mutation in ORF50 that leads to a stop codon at amino acid 116 in RTA. Nucleotides 2,022 to 9,739 were subsequently removed to generate RDV-50.stopΔM1-M4 (Supplementary Figure 1a). The ΔM1-M4 deletion was validated by sequencing and restriction fragment length polymorphism analysis, confirming the expected change upon *Hind*III digestion (Supplementary Figure 1a,b).

To analyze the virus-specific immune response generated by the administration of RDV-50.stopΔM1-M4 in comparison to RDV-50.stop, WT C57BL/6 mice were inoculated intraperitoneally (IP) with 1X10^6^ PFU of either RDV-50.stopΔM1-M4 or RDV-50.stop or sham vaccinated at d0 for the prime vaccination and again at d28 for the boost vaccination (Figure 1a). At d28 post-boost, CD8+ T cells responses in lungs and spleen were evaluated by flow cytometry for activation (CD44^hi^) and virus specificity by binding to MHC-I complexes presenting the p56 immunodominant epitope derived from ORF6 or the p79 immunodominant epitope derived from the ORF61 ribonucleotide reductase large subunit [44]. In comparison to naïve animals, CD8 T cells specific to MHV68 p56 epitopes were detected at higher levels in lungs of vaccinated mice (Figure 1b,c). Levels of p56 virus-specific T cells were comparable in mice vaccinated with either RDV-50.stopΔM1-M or RDV-50.stop (Figure 1b,c).

**Figure 1.**
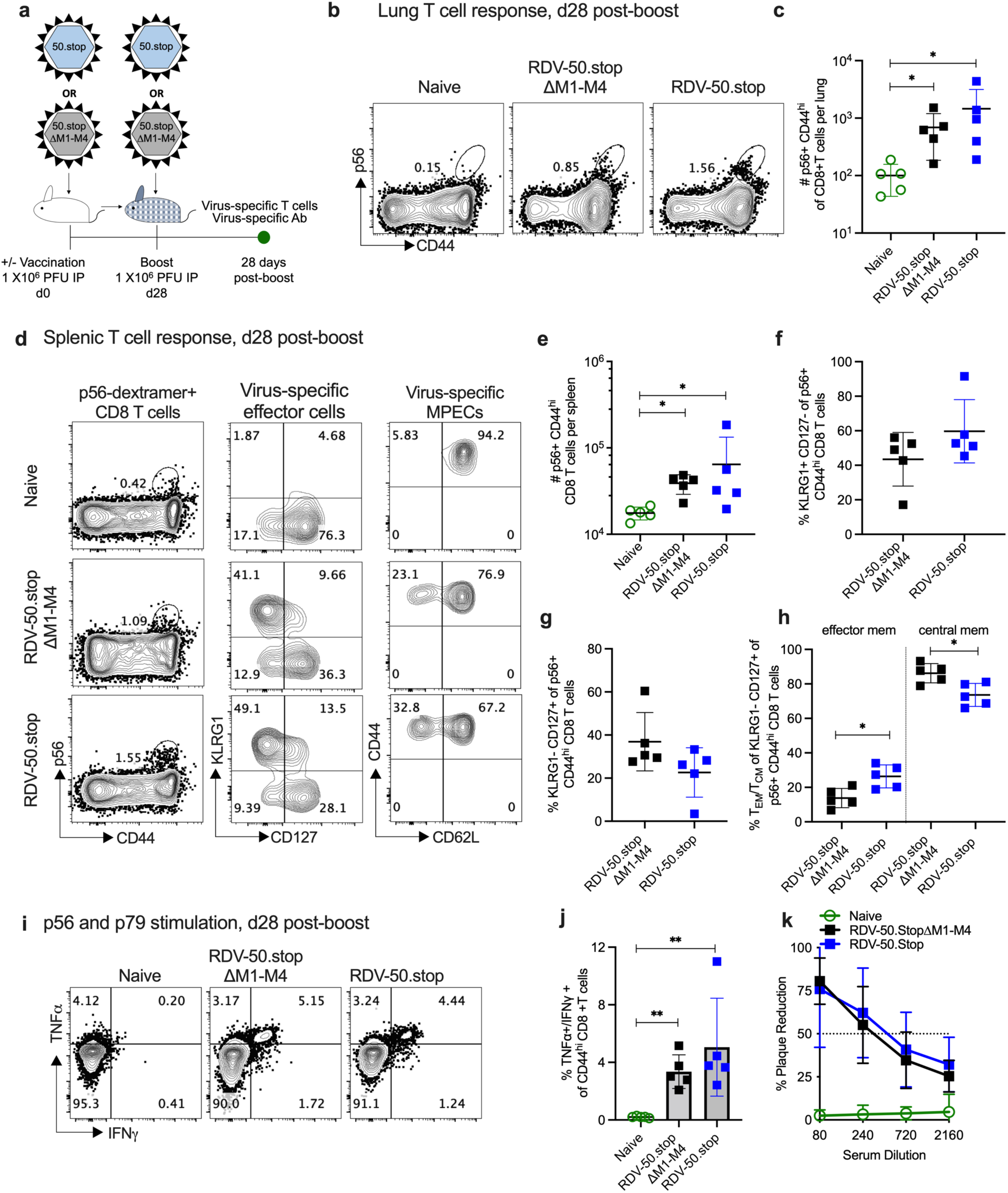
RDV-50.stopΔM1-M4 MHV68 vaccine generates virus-specific immune responses upon a prime-boost regimen in C57BL/6 mice. (**a**) Schematic of prime-boost strategy to examine the immune response of C57BL/6 mice to RDV-50.stopΔM1-M4 or RDV-50.stop vaccination. C57BL/6 mice were vaccinated IP twice (prime+boost) with 1×10^6^ PFU RDV-50.stopΔM1-M4 or RDV-50.stop. Naïve mice were age-matched, non-vaccinated controls. (**b**) Flow cytometric gating strategy to determine the frequency of CD44^hi^ CD8 T cells in the lungs that were reactive with the MHV68 p56 epitope at d28 post-boost. (**c**) Total p56-dextramer+ CD8 T cells per lung of individual mice. (**d**) Left column, flow cytometric gating strategy to determine the frequency of CD44^hi^ CD8 T cells in the spleens that were reactive with viral p56. Middle column, p56-dextramer+ CD8 T cells were further analyzed for markers of short-lived effector cell (SLEC, KLRG1^+^CD127^-^) and memory precursor effector cell (MPEC, KLRG1^-^CD127^+^) subsets. Right column, MPECs were further delineated into CD62L^-^ effector and CD62L^+^ central MPECs. (**e**) Total p56-dextramer+ CD8 T cells per spleen of individual mice with the indicated vaccination regimen. The percentage of p56-specific CD8 T cells that were SLECs (**f**), MPECs (**g**), and effector vs memory MPECs (**h**) were enumerated based on the gating strategy for surface markers in (d). (**i**) Flow cytometric gating strategy to determine the intracellular cytokine levels of effector cytokines TNFα and IFNγ at 5 hrs after dual stimulation with p79 and p56 peptides. (**j**) Percentage of CD44^hi^ CD8 T cells producing both TNFα and IFNγ. (**k**) Virus neutralization in serum of vaccinated mice at d28 post-boost determined by a plaque reduction assay. For (c, e-h, j, k) symbols represent individual mice (N=5). Bars and whiskers are mean +/- SD. *, p<0.05; **, p<0.01 in Sidak’s (c) or Kruskal-Wallis (e) multiple comparisons test of one-way ANOVA and in two-tailed unpaired t test (f-h) and Mann-Whitney test (j) between the indicated groups.

RDV-50.stopΔM1-M4 vaccination induced abundant virus-specific CD8 T cells in the spleens that were comparable to RDV-50.stop vaccination (Figure 1 d-h; Supplementary Figure 2a-d). Splenic CD8 T cells were further identified as KLRG1+CD127-short-lived effector cells (SLECs) and KLRG1-CD127+ memory precursor effector cells (MPECs). Virus-specific SLECs (Figure 1d,f; Supplementary Figure 2b) and MPECs (Figure 1d,g; Supplementary Figure 2c) were comparable in mice vaccinated with either RDV-50.stopΔM1-M or RDV-50.stop. Vaccination with RDV-50.stopΔM1-M4 yielded modest changes in memory MPEC populations.RDV-50.stop elicited more CD62L-effector memory MPEC and RDV-50.stopΔM1-M4 elicited more CD62L+ central memory MPEC (Figure 1d,h; Supplementary Figure 2d). For both vaccines, stimulation of splenic CD8 T cells with a combination of p56 and p79 peptides led to increased production of antiviral effector cytokines TNFα and IFNγ in vaccinated mice compared to naïve mice (Figure 1i,j).

The neutralization capacity of serum from vaccinated mice was determined by plaque reduction neutralization assays (PRNT; Figure 1k). PRNT50 titers, the dilution of serum to reach 50% neutralization of plaques, indicated that RDV-50.stopΔM1-M4 elicited comparable levels of virus-neutralizing antibodies to RDV-50.stop. The percentage of Vβ4+ CD8 T cells in pooled spleens of prime-boost vaccinated mice and WT MHV68 infected mice at d28 post inoculation was analyzed by flow cytometry. As expected, Vβ4+ CD8 T cells were increased upon M1 expression in WT MHV68 infection (Supplementary Figure 2e), but were not induced upon vaccination with either RDV-50.stopΔM1-M4 that lacks M1 or RDV-50.stop that do not express M1 transcript [23]. Taken together, RDV-50.stopΔM1-M4 vaccination elicits virus-specific CD8 T cell responses and neutralizing antibodies that are comparable to RDV-50.stop.

### Vaccination with RDV-50.stopΔM1-M4 protects mice from WT MHV68 challenge

To determine if the immunity generated by RDV-50.stopΔM1-M4 vaccination was protective *in vivo*, prime-boost vaccinated C57BL/6 mice were challenged by IN infection with 1000 PFU of WT MHV68 at d28 post-boost and compared to naïve and RDV-50.stop vaccinated animals (Figure 2a). Plaque assays performed on homogenized lung tissue at d7 post-challenge revealed significantly lower viral titers in lungs of vaccinated animals compared to sham-vaccinated controls (Figure 2b). Lung titers were also analyzed at d7 post WT challenge with a higher 5000 PFU dose. In this case, virus replication in the lungs was detected in 2 of 5 RDV-50.stopΔM1-M4 vaccinated mice and in 4 of 5 RDV-50.stop vaccinated mice upon challenge, but there was a significant ∼2.5 log reduction in viral titers upon vaccination with either RDV-50.stopΔM1-M4 or RDV-50.stop compared to sham-vaccinated controls (Supplementary Figure 3a). These data demonstrate that vaccination with RDV-50.stopΔM1-M4 markedly reduces WT virus replication at a mucosal barrier.

**Figure 2.**
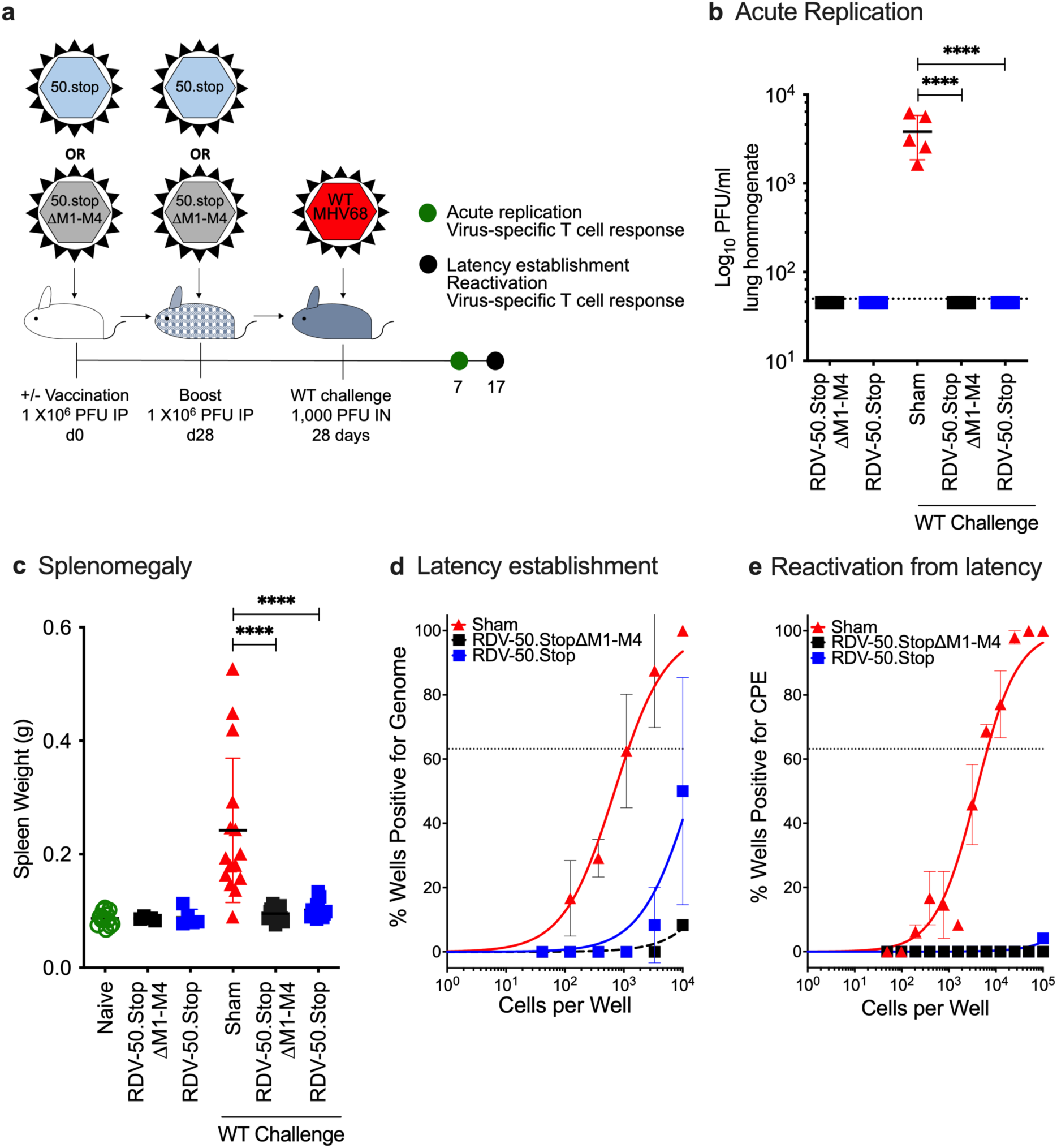
Vaccination with RDV-50.stopΔM1-M4 reduces acute replication, splenomegaly, latency and reactivation of wild-type MHV68 upon challenge in C57BL/6 mice. (**a**) Schematic describing C57BL/6 mice either sham vaccinated or prime-boost vaccinated IP with 1×10^6^ PFU of RDV-50.stopΔM1-M4 or RDV-50.stop followed by IN challenge with 1×10^3^ PFU of WT MHV68 at d28 post-boost. Naïve mice were age-matched, non-vaccinated controls. (**b**) Acute replication at d7 post-challenge determined by measuring infectious particles per ml lung homogenate. (**c**) Splenomegaly determined by spleen weights at d17 post-challenge. Symbols denote individual mice (N=5-15 mice per experiment); bars and whiskers represent mean +/- SD. ****, p<0.0001 in Sidak’s multiple comparisons test of one-way ANOVA (**b, c**) between the indicated groups. (**d**) The frequency of latency determined by limiting dilution nested PCR of intact splenocytes for the viral genome at d17 post-challenge. (**e**) The frequency of explant reactivation determined by limiting dilution coculture of intact viable splenocytes on a monolayer of primary MEFs d17 post-challenge. Disrupted splenocytes plated in parallel did not detect preformed infectious virus in the vaccinated animals. For (**d-e**), symbols denote the average of two experiments with five mice per experiment; error bars represent SEM.

After acute replication in the lung tissue, MHV68 transits the draining lymph nodes and disseminates to the spleen via the bloodstream [10]. An increase in spleen weight, termed splenomegaly, is akin to infectious mononucleosis caused by EBV and coincides with virus colonization of the spleen [45,46]. WT challenge of sham-vaccinated animals led to a mean 2.8-fold increase in splenomegaly at d17 post-challenge (Figure 2c). Vaccination with either RDV-50.stopΔM1-M4 or RDV-50.stop alone did not cause an increase in spleen weight. Vaccination with either RDV-50.stop prior to WT challenge prevented infection-induced splenomegaly (Figure 2c). Limiting-dilution PCR analyses were performed to measure the frequency of splenocytes that harbor MHV68 genomes. Low frequencies of viral-genome positive splenocytes were detected upon RDV-50.stop vaccination alone, but not in mice vaccinated with RDV-50.stopΔM1-M4 (Supplementary Figure 3b). Consistent with the loss of RTA encoded by ORF50, vaccination with RDV-50.stopΔM1-M4 or RDV-50.stop did not result in reactivation from splenocytes (Supplementary Figure 3c). With regard to protection, vaccination prior to WT virus challenge resulted in a significant reduction in latently infected cells from 1 in 1149 splenocytes in sham-vaccinated mice to levels below the limit of detection for either RDV-50.stop vaccine (Figure 2d). Consistent with the significant reduction in splenic latency, vaccination with either RDV-50.stop resulted in undetectable reactivation in a limiting-dilution *ex vivo* evaluation of cytopathic effects on an indicator monolayer (Figure 2e). In contrast, splenocytes from sham-vaccinated mice exhibited a typical *ex vivo* reactivation frequency of 1 in 6497 splenocytes. These data indicate that RDV-50.stopΔM1-M4 vaccination is as effective as RDV-50.stop in blocking both latency establishment and the subsequent reactivation from latency at d17 post WT challenge, the peak time of MHV68 splenic colonization in mice.

### Virus-specific immune responses following vaccination with RDV-50.stopΔM1-M4 correlate with protection from WT challenge

To develop an effecive GHV vaccine, it is important to identify the correlates of immune protection. We therefore evaluated virus-specific immune responses following WT challenge of mice vaccinated with RDV-50.stopΔM1-M4 or RDV-50.stop. At d7 post WT MHV68 challenge, CD8 T cell responses against the p56 immunodominant epitope were significantly higher in lungs of vaccinated mice in comparison to sham-vaccinated controls (Figure 3a). Stimulation of splenocytes with p56 and p79 peptides led to increased production of antiviral effector cytokines TNFα and IFNγ in both cohorts of RDV vaccinated mice (Figure 3b) compared to sham-vaccinated mice, demonstrating that antigen-specific cells were present and responsive. Total p56-specific CD8 T cells were slightly elevated in spleens of vaccinated animals (Figure 3c), and total p79-specific CD8 T cells were significantly higher in either vaccinated group compared to sham-vaccinated controls (Figure 3g). Further immunophenotyping revealed that over 30% of p56-specific (Figure 3d) and 50% of p79-specific (Figure 3h) CD44^hi^ CD8 T cells in vaccinated mice were SLECs. As expected, mice vaccinated with either RDV-50.stop had higher levels of SLECs than MPECS compared to sham-vaccinated mice (Figure 3d,e,h,i). CD62L analysis of MPECs identified a similar proportion of CD62L− effector memory MPEC compared to CD62L+ central memory MPEC in both vaccinated groups compared to sham-vaccinated mice (Figure 3f,j). Taken together, prime-boost vaccination with RDV-50.stopΔM1-M4 elicited virus-specific CD8 T cells that were comparable to RDV-50.stop vaccination. Thus, virus-specific CD8 T effector responses correlate with strong protection afforded to RDV-50.stop vaccinated mice against WT MHV68 infection.

**Figure 3.**
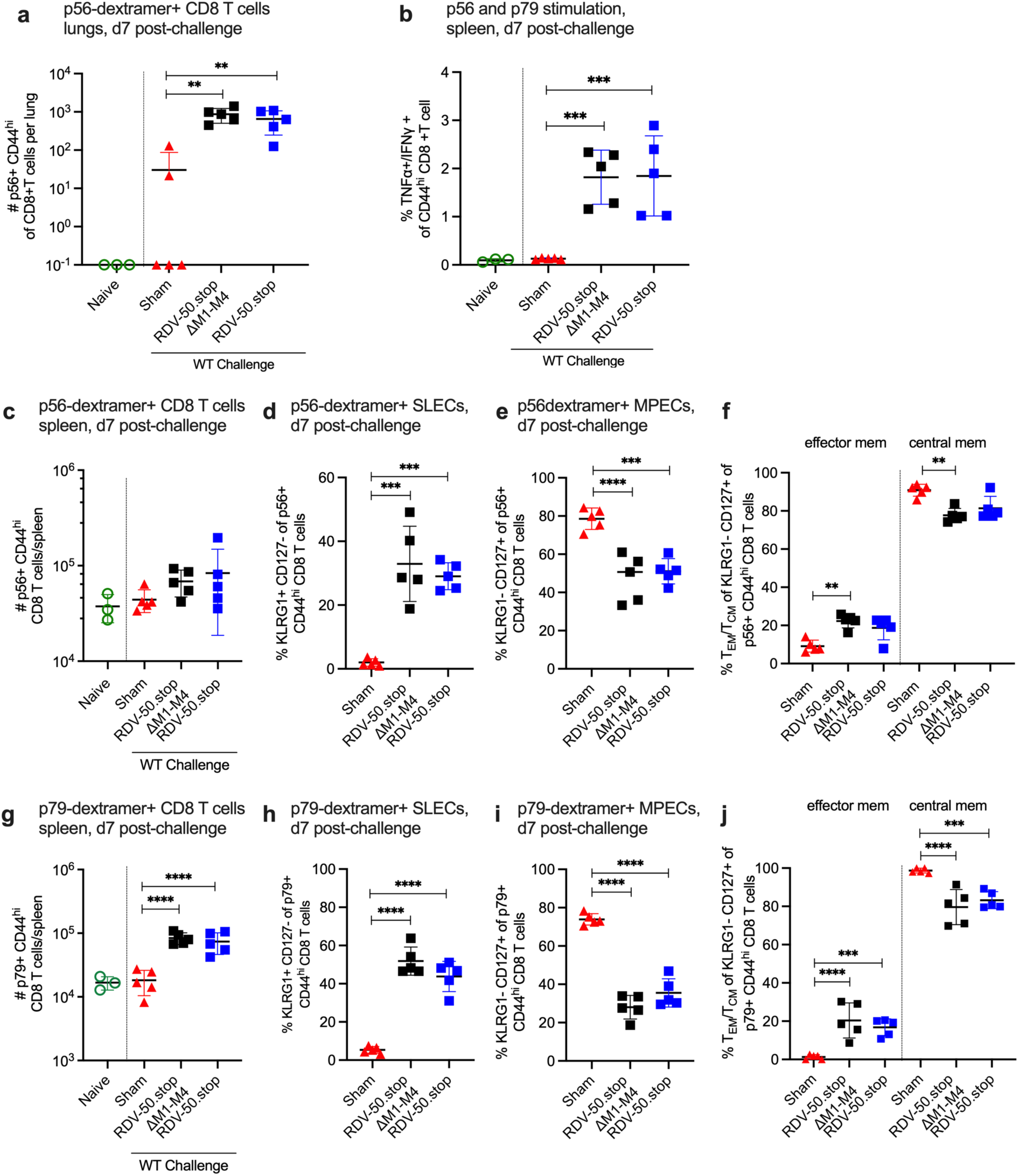
Evaluation of T cell response to MHV68 in the lungs and spleens of vaccinated mice at seven days post-challenge with WT virus. C57BL/6 mice were either sham vaccinated or prime-boost vaccinated IP with 1×10^6^ PFU RDV-50.stopΔM1-M4 or RDV-50.stop followed by IN challenge with 1×10^3^ PFU WT MHV68 at d28 post-boost and analyzed d7 post-challenge. **(a)** Total p56-dextramer+ CD8 T cells per lung of individual mice after prime+boost+challenge. **(b)** Intracellular cytokine levels of effector cytokines TNFα and IFNγ were examined 6 hrs after dual stimulation with p56 and p79 peptides. Total **(c)** p56-dextramer+ or **(g)** p79-tetramer+ CD8 T cells per spleen of individual mice with the indicated vaccination regimen. Percentage of p56-tetramer+ or p79-dextramer+ CD8 T cells **(d,h)** with markers of short-lived effector cell (SLEC, KLRG1^+^CD127^-^) and **(e,i)** memory precursor effector cell subsets (MPEC, KLRG1^-^CD127^+^). (**f,h)** MPECs were further delineated into CD62L^-^ effector and CD62L^+^ central MPECs for p56-dextramer+ or p79-dextramer+ CD8 T cells. For each graph, symbols represent individual mice, (N=3-5); bars and whiskers are mean +/- SD. **, p<0.01; ***, p<0.001; ****, p<0.0001 in Sidak’s multiple comparisons test of one-way ANOVA between the indicated groups.

We also evaluated the effect of RDV-50.stopΔM1-M4 vaccination on virus-specific CD8 T cell responses to WT virus challenge on d17 post-challenge, a timepoint that approximates the typical peak of latency establishment in the spleen after IN infection. The quantity of virus-specific CD8 T cells observed at d17 post-challenge in the lungs of sham-vaccinated mice surpassed the levels detected in the RDV-50.stop vaccinated mice upon challenge (Figure 4a). In the case of the spleen at d17 post-challenge, p56-specific CD8 T cells were not more numerous in sham vaccinated mice compared to RDV-50.stopΔM1-M4 vaccinated mice, but exhibited an increased SLEC and CD62L-effector MPEC phenotype (Figure 4b,c,e). The p79-specific CD8T cells were more abundant in sham vaccinated mice with a similar effector cell phenotype compared to vaccinated mice (Supplementary Figure 4a,b,d). In summary, the virus-specific CD8 T cell response in both RDV-50.stopΔM1-M4 and RDV-50.stop vaccinated mice was dampened compared to sham-vaccinated mice at d17 post-challenge. This observation is consistent with a strong pre-existing immune response in the lungs and spleen elicited by RDV-50.stop vaccination that effectively controlled virus replication at d7 (Figure 2). In contrast, sham-vaccinated mice that are unable to control initial virus replication mount a stronger effector CD8 T cell response at d17 post-challenge.

**Figure 4.**
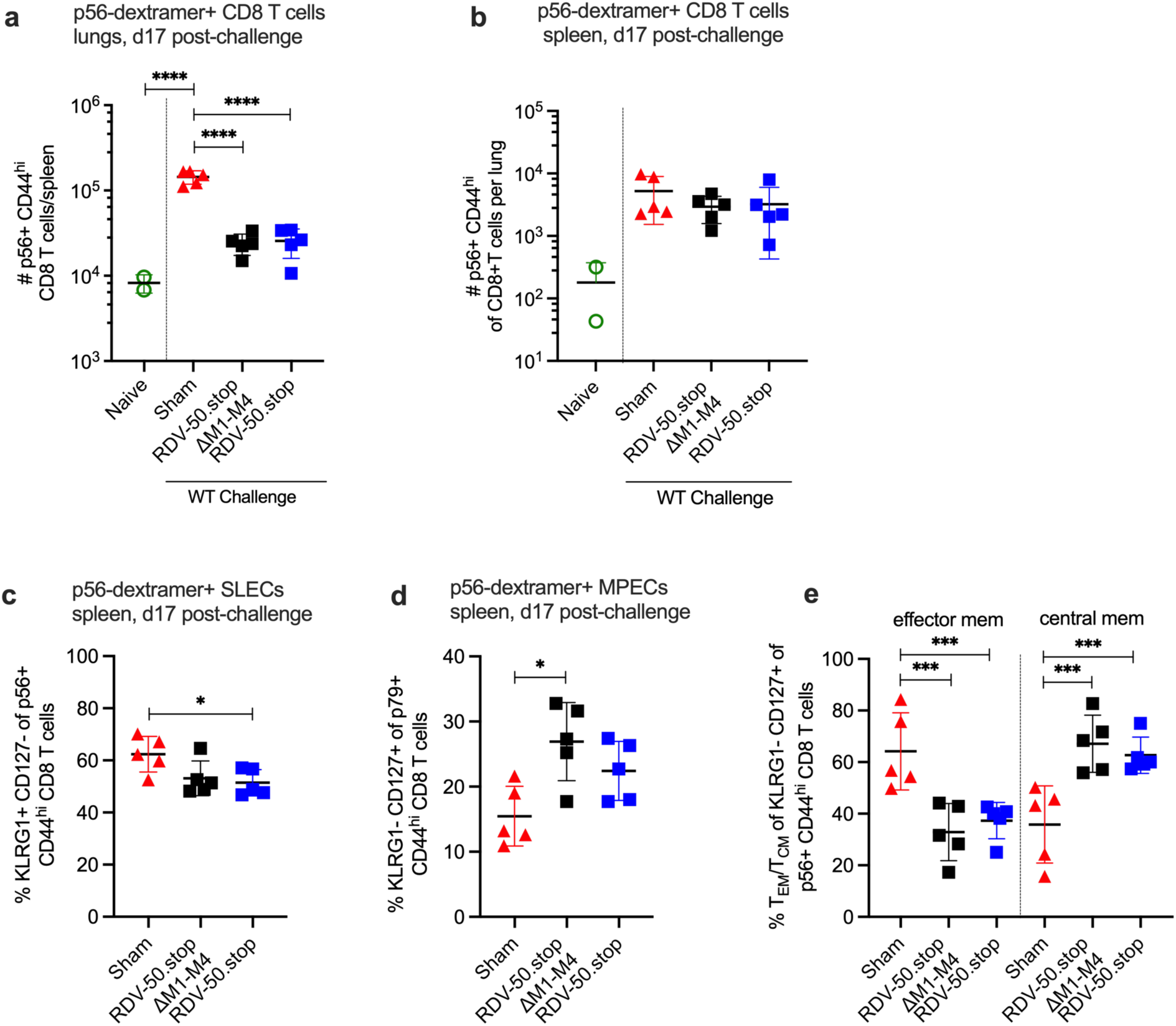
Evaluation of T cell response to MHV68 in the lungs and spleens of vaccinated mice at seventeen days post-challenge with WT virus. C57BL/6 mice were either sham-vaccinated or prime-boost vaccinated IP with 1×10^6^ PFU RDV-50.stopΔM1-M4 or RDV-50.stop followed by challenge with 1×10^3^ PFU WT MHV68 at d28 post-boost and analyzed d17 post-challenge. (**a**) Total p56-dextramer+ CD8 T cells per lung of individual mice after prime+boost+challenge. (**b**) Total p56-dextramer+ CD8 T cells per spleen of individual mice with the indicated vaccination regimen. Percentage of p56-dextramer+ CD8 T cells with markers of (**c**) short-lived effector cell (SLEC, KLRG1^+^CD127^-^) and (**d**) memory precursor effector cell (MPEC, KLRG1^-^CD127^+^) subsets. (**e**) MPECs were further delineated into CD62L^-^ effector and CD62L^+^ central MPECs for p56-dextramer+ CD8 T cells. For each graph, symbols represent individual mice, (N=2-5); bars and whiskers are mean +/- SD. *, p<0.05; ***, p<0.001; ****, p<0.0001 in Sidak’s multiple comparisons test of one-way ANOVA between the indicated groups.

## Discussion

The mouse gammaherpesvirus model is critical to inform vaccine design for KSHV and EBV and to identify the immune components that mediate protection against GHV in the host. In this study, we further refined the RDV-50.stop vaccine platform with the deletion of major pathogenesis determinants of MHV68 *in vivo* that includes the M1-M4 genes and non-coding TMERs 6-8. We report that prime-boost vaccination with RDV-50.stopΔM1-M4 elicits virus-neutralizing antibodies and virus-specific CD8 T-cell responses in lungs and spleen that were comparable to RDV-50.stop vaccination. Importantly, mice vaccinated with either RDV-50.stop were protected from acute virus replication in lungs, and exhibited a reduction in latency establishment coupled with near-complete abolishment of virus reactivation in splenocytes. Taken together, several pathogenic determinants unique to MHV68 are dispensable for eliciting a protective immune response.

At the onset of our study, the net impact of the loss of the M1-M4 latency locus on the host immune response to RDV-50.stop had not been reported. M1 drives Vβ4+ CD8 T cell expansion that contributes to control of reactivation from latency [26]. Vβ4+ CD8 T cells were not induced in RDV-50.stopΔM1-M4 vaccinated mice, as expected. Given that RDV-50.stopΔM1-M4 vaccination was still effective at protection from WT challenge, we conclude that Vβ4+ CD8 T cells are not an essential immune component for the protection of C57BL/6 mice against the early phases of MHV68 infection in the lung and splenic tissues. Similarly, loss of M2, a known source of cytotoxic T cell epitopes [30,31], did not negatively impact the elicitation of protective responses in vaccinated animals. On the other hand, the M3 gene encodes a chemokine-binding protein that dampens immune responses by inhibiting both antiviral and inflammatory activities during infection [32–35]. Given that inflammation may promote immune recruitment and activation, eliminating the M3 gene in a vaccine candidate could potentially improve immunogenicity. However, we observed no enhanced immune response when vaccinating in the absence of the M1-M4 latency locus. Our results are consistent with findings from another study that reported the removal of M1, M2, M3, M4, and ORF4 had no impact on protection afforded by vaccination with an MHV68 mutant lacking ORF73 which encodes for the latency associated nuclear antigen [47].

In our previous study of RDV-50.stop vaccination [23], we observed low frequencies of splenocytes harboring the RDV-50.stop genome, indicating that it could establish a persistent infection even though reactivation was not possible. This is not an ideal endpoint for a GHV vaccine that expresses viral genes with oncogenic potential. The M2 and M4 genes each contribute to the establishment of latency in the B cell reservoir by MHV68, leading to the hypothesis that their absence would further attenuate latency. Here, mice vaccinated with RDV-50.stopΔM1-M4 had qualitatively lower levels of genome-positive cells compared to those vaccinated with RDV-50.stop, albeit the frequency of splenocytes that harbor either vaccine-derived genome were reduced to levels below the limit of absolute quantitation.

We previously reported that RDV-50.stop vaccinated mice were protected from WT virus replication in the lungs [23]. In this study, we demonstrated that both RDV-50.stopΔM1-M4 and RDV-50.stop protect against acute replication upon IN challenged with 1000 PFU WT MHV68. However, when vaccinated mice were challenged with an increased dose of 5000 PFU, there was a higher breakthrough in acute replication in the lungs. This dose-dependent incidence of breakthrough infection indicates that mucosal immune defenses might be further improved via additional adjuvanted boosts.

Taken together, four ORFs and three TMERs that are unique to MHV68 and support the pathogenesis of WT virus are not required to elicit protective immunity in the context of RDV-50.stop vaccination. We conclude that RDV-50.stopΔM1-M4 is appropriate to further develop as a safe and effective vaccine platform in the mouse GHV pathogen system.

## Patents

JCF and LTK hold a patent [US Patent 11,149,255] for the design and production of the codon-shuffled RTA (CS-RTA)-expressing producer cells to prevent WT revertant of mutants. LTK is divested from potential earnings.

## Supplementary Materials

Table S1: Primers, gBlocks and antibodies used in this study. Figure S1: Generation of recombinant RDV-50.stopΔM1-M4 MHV68 upon deletion of unique M gene locus.; Figure S2: RDV-50.stopΔM1-M4 generates virus-specific immune responses upon a prime-boost regimen in C57BL/6 mice.; Figure S3: Vaccination with RDV-50.stopΔM1-M4 reduces acute replication, latency and reactivation in C57BL/6 mice.; Figure S4: Evaluation of T cell response to MHV68 in the spleens of vaccinated mice at seventeen days post-challenge with WT virus.

## Author Contributions

Conceptualization, J.C.F., L.T.K.; methodology, D.M., D.O., L.T.K; validation, D.M., D.O.; formal analysis, D.M.; investigation, D.M.,D.O., L.T.K.; resources, D.O., J.C.F., L.T.K.,; data curation, D.M.; writing—original draft preparation, D.M., D.O., L.T.K.; writing—review and editing, D.M., D.O., J.C.F., L.T.K.; visualization, D.M., D.O., L.T.K.; supervision, L.T.K.; project administration, J.C.F., L.T.K.; funding acquisition, J.C.F., L.T.K. All authors have read and agreed to the published version of the manuscript.

## Funding

This research was supported in part by the Intramural Research Programs of the NIH, the National Cancer Institute (ZIA BC 012014) (L.T.K., D.M.). J.C.F. was supported by NIH R01CA167065 and R01AI181787.

## Institutional Review Board Statement

All animal procedures reported in this study were performed by NCI-CCR affiliated staff, approved by the NCI Animal Care and Use Committee (ACUC) and in accordance with federal regulatory requirements and standards. All components of the intramural NIH ACU program are accredited by AAALAC International.

Mice were anesthetized prior to inoculation. Mice were humanly euthanized at the indicated experimental endpoints. Humane endpoint criteria such as signs of distress including lethargy, dehydration, or a bodyweight reduction of 20% were not reached.

## Acknowledgments

The authors thank members of the Krug laboratory for the valuable feedback and insightful discussions.

## Conflicts of Interest

The authors declare no conflicts of interest.

## Notes

### Competing Interest Statement

The authors have declared no competing interest.

