## Supplemental Files for "Vaccination with a Replication-Dead Murine Gammaherpesvirus Lacking Viral Pathogenesis Genes Inhibits WT Virus Infection"

Supplemental Materials

Supplemental Methods

*Restriction fragment length polymorphism:* To confirm the integrity of the MHV68 BACmid after genetic manipulation, RFLP analysis was performed by *Hind*III restriction enzyme digestion of 1ug of BACmid DNA. Clone# 121 (RDV-50.stop), clone# 466 (RDV-50.stopΔM1-M4), and WT MHV68 BAC by overnight 37° C. Fragments were separated by 0.9% agarose electrophoresis at 30V for 20 hours.

*Vβ4+ CD8 T cells analysis by flow cytometry:* Spleens were processed using a tissue grinder and were treated to remove RBC and filtered as described previously prior to resuspension in 200 µl of PBS with 2% fetal bovine serum and blocked with TruStain fcX (BioLegend, San Diego, CA). T cell subsets were identified with antibodies against CD45 (dilution 1:200; clone 30-F11; BV510; cat# 563891), Vβ4+ (dilution 1:100; clone KT4; FITC; cat# 553365) purchased from BD Biosciences, Franklin Lakes, NJ and CD4 (dilution 1:400; clone GK1.5; PerCP-Cy5.5; cat# 100434), CD8 (dilution 1:400; clone 53-6.7; Pacific Blue; cat# 100725), CD62L (dilution 1:2000; clone MEL-14; PE; cat# 104408), CD44 (dilution 1:1000; clone IM7; APC; cat# 103012) purchased from BioLegend, San Diego, CA. The data was collected on a CytoFLEX flow cytometer (Beckman Coulter, Brea, CA) and analyzed using FlowJoX v10.10.0 (Treestar Inc., Ashland, OR). CD8+ T cells were first gated as lymphocytes based on forward and side scatter parameters, singlets, then live per exclusion of Alexa Fluor™ 700 NHS Ester uptake, followed by gating for Vβ4+ CD8+CD45+.

| Table S1. Primers, gBlocks and antibodies used in this study | |
|---|---|
| **Mutant MHV68 generation** | |
| **Mutation** | **gBlock[a]** |
| ORF50 locus<br>50_STOP_FS_gBLK_1 | AGAAACCAGAAGGTGAGGTTTAATGCCAAAGTCCATAACAGGCATCCATGTGGGTACATA**TAG**TCTC<br>ACCACCTGATC**T**AAATATGCCATTGA**T**AAGAGTTGTCTA**G**ACCACAGACAGGCTGTTTCTTAGGGATA<br>ACAGGGTAATCGATTTATTCAACAAAGCCACGTTGTGTCTCAAAATCTCTGATGTTACATTGCACAAGA<br>TAAAAATATATCATCATGAACAATAAAACTGTCTGCTTACATAAACAGTAATACAAGGGGTGTTATGA<br>GCCATATTCAACGGGAAACGTCTTGCTCGAGGCCGCGATTAAATTCCAACATGGATGCTGATTTATATG<br>GGTATAAATGGGCTCGCGATAATGTCGGGCAATCAGGTGCGACAATCTATCGATTGTATGGGAAGCCC<br>GATGCGCCAGAGTTGTTTCTGAAACATGGCAAAGGTAGCGTTGCCAATGATGTTACAGATGAGATGGT<br>CAGACTAAACTGGCTGACGGAATTTATGCCTCTTCCGACCATCAAGCATTTTATCCGTACTCCTGATGA<br>TGCATGGTTACTCACCACTG |
| ORF50 locus<br>50_STOP_FS_gBLK_2 | CTCCTGATGATGCATGGTTACTCACCACTGCGATCCCCGGGAAAACAGCATTCCAGGTATTAGAAGAA<br>TATCCTGATTCAGGTGAAAATATTGTTGATGCGCTGGCAGTGTTCCTGCGCCGGTTGCATTCGATTCCTG<br>TTTGTAATTGTCCTTTTAACAGCGATCGCGTATTTCGTCTCGCTCAGGCGCAATCACGAATGAATAACG<br>GTTTGGTTGATGCGAGTGATTTTGATGACGAGCGTAATGGCTGGCCTGTTGAACAAGTCTGGAAAGAA<br>ATGCATAAGCTTTTGCCATTCTCACCGGATTCAGTCGTCACTCATGGTGATTTCTCACTTGATAACCTTA<br>TTTTTGACGAGGGGAAATTAATAGGTTGTATTGATGTTGGACGAGTCGGAATCGCAGACCGATACCAG<br>GATCTTGCCATCCTATGGAACTGCCTCGGTGAGTTTTCTCCTTCATTACAGAAACGGCTTTTTCAAAAAT<br>ATGGTATTGATAATCCTGATATGAATAAATTGCAGTTTCATTTGATGCTCGATGAGTTTTTCTAATCAGA<br>ATTGGTTAATTGGTTGTAACACTGGCGGCATCCATGTGGGTACATA**TAG**TCTCACCACCTGATC**T**AAAT<br>ATGCCATTGA**T**AAGAGTTGTCTA**G**ACCACAGACAGGCTGTTTCTGACTGCCCCATGTTCGGGGGTACA<br>GCTACCTCAACCTCTG |
| M1-M4 deletion<br>M1_4DEL gBLK_1 | CCTTTATTGTAAGGGTACTCTCATCACCAATGTAAATTAATATGTAGCAAACTTTGGTGTGGGAGTCCT<br>ACCCCTTTTGCTCCACAGGCCGCCCACGACCATCTAGCCTCCCACCAAAACAAGAACAGTTGCAGCTT<br>TTGCTGTTTTCCTTAATATTTATCTTTGTGGTGTTCAGTCAGGCCTTGCTCTCAGTTTCTATGCCCCCAGG<br>CTGGTTTCAAACACTGGTCAGAAGGCTATCTTTCTTGGTGGTTTCACTTCTAAACATGGGCCATTAAAA<br>GGGAGGGAATTGGCATCATTGAGCAGCGGCGACCTAGGGATAACAGGGTAATCGATTTATTCAACAA<br>AGCCACGTTGTGTCTCAAAATCTCTGATGTTACATTGCACAAGATAAAAATATATCATCATGAACAAT<br>AAAACTGTCTGCTTACATAAACAGTAATACAAGGGGTGTTATGAGCCATATTCAACGGGAAACGTCTT<br>GCTCGAGGCCGCGATTAAATTCCAACATGGATGCTGATTTATATGGGTATAAATGGGCTCGCGATAAT<br>GTCGGGCAATCAGGTGCGACAATCTATCGATTGTATGGGAAGCCCGATGCGCCAGAGTTGTTTCTGAA |

| | |
|---|---|
| | ACATGGCAAAGGTAGCGTTGCCAATGATGTTACAGATGAGATGGTCAGACTAAACTGGCTGACGGAA TTTATGCCTCTTCCGACCATCAAGCATTTTATCCGTACTCCTGATGATGCATGGTTACTCACCACTG |
| M1-M4 deletion<br>M1_4DEL gBLK_2 | CTCCTGATGATGCATGGTTACTCACCACTGCGATCCCCGGGAAAACAGCATTCCAGGTATTAGAAGAA TATCCTGATTCAGGTGAAAATATTGTTGATGCGCTGGCAGTGTTCCTGCGCCGGTTGCATTCGATTCCTG TTTGTAATTGTCCTTTTAACAGCGATCGCGTATTTCGTCTCGCTCAGGCGCAATCACGAATGAATAACG GTTTGGTTGATGCGAGTGATTTTGATGACGAGCGTAATGGCTGGCCTGTTGAACAAGTCTGGAAAGAA ATGCATAAGCTTTTGCCATTCTCACCGGATTCAGTCGTCACTCATGGTGATTTCTCACTTGATAACCTTA TTTTTGACGAGGGGAAATTAATAGGTTGTATTGATGTTGGACGAGTCGGAATCGCAGACCGATACCAG GATCTTGCCATCCTATGGAACTGCCTCGGTGAGTTTTCTCCTTCATTACAGAAACGGCTTTTTCAAAAAT ATGGTATTGATAATCCTGATATGAATAAATTGCAGTTTCATTTGATGCTCGATGAGTTTTTCTAATCAGA ATTGGTTAATTGGTTGTAACACTGGCTTCTAAACATGGGCCATTAAAAGGGAGGGAATTGGCATCATT GAGCAGCGGCGACCTTACATTCATCTGGGAATATGGTATTGAGATTTATGACTTTCTAGAATAACTGTA CCCTGTTAAGTTTCAATTCCTTGTGGCCCTACCCCGAATCTCTATTAAAGGGTTAATAAAAATTACTCTC AACAAAATCATGGCCAACTTCCACTTTTTCTGCGCAGTATTGGTGGGGATTGTGGGTGTAAATGGTGAC AACATGTGCCACCATCTTCCTCAAATGCCCACACTAACATCTCCTAAATTTACACCACCAGTCAAAAG TGGCACCACCCTGCAGCTCCGCTGTAGGCCAGGGTTCACACCAGGCGCA |
| **Mutation** | **Amplification Primers** |
| RDV-50.stop<br>50_STOP_FS_FOR | 5′AGAAACCAGAAGGTGAGGTTTAATG |
| RDV-50.stop<br>50_STOP_FS_REV | 5′CAGAGGTTGAGGTAGCTGTACCC |
| RDV-50.stopΔM1-M4<br>M1-4_DEL_FOR | 5′-CACCAATGTAAATTAATATGTAGC |
| RDV-50.stopΔM1-M4<br>M1-4_DEL_REV | 5′-AACCCTGGCCTACAGCGGAGCTGC |
| **Mutation** | **Sanger Sequencing Primers** |
| ORF50 locus<br>50_FSS_SEQ_FOR | 5′ACAAATTTTACACAGCACCTGAAGC |
| ORF50 locus<br>50_FSS_SEQ_REV | 5′ATGCCTCAACTTCTCTGGATATG |

| | |
|---|---|
| <u>M1-M4 deletion</u><br>M1-4_SEQ_FOR | 5′-CACCAATGTAAATTAATATGTAGC |
| <u>M1-M4 deletion</u><br>M1-4_SEQ_REV | 5'-GGTGGTGGTGCTCATGTCTGACG |

[a]Bold letters - **TAG** and two **T’s** were added, then a **G** was substituted for a T to obtain a *Xba*I site for screening which sufficiently disrupts the reading frame. Underlined-frameshift region

Supplemental Figure Legends

**Supplementary Figure 1 Generation of recombinant RDV-50.stopΔM1-M4 MHV68 upon deletion of unique M gene locus. (a)** Schematic describing the left end of WT MHV68 and the parental RDV-50.stop. M1- M4 genes and TMERs 6-8 (nucleotides 2022 to 9739) were deleted from RDV-50.stop to generate MHV68 RDV-50.stopΔM1-M4. '*' and '^' denote HindIII digestion sites. '*' indicates the site lost with the deletion of 2022 to 9739 bp. Figure made with BioRender. **(b)** In the HindIII restriction digest, WT MHV68 and RDV-50.stop has the expected 6156 bp fragment, while the RDV-50.stopΔM1-M4 genome has the expected loss of the 6156 bp fragment and gain of a 3277 bp fragment, indicated by red arrows.

**Supplementary Figure 2 RDV-50.stopΔM1-M4 generates virus-specific CD8 T cell responses but does not induce M1 driven Vβ4+ CD8 T cells upon a prime-boost regimen in C57BL/6 mice.** C57BL/6 mice were either sham-vaccinated or prime-boost vaccinated IP with $1x10^6$ PFU RDV-50.stopΔM1-M4 or RDV-50.stop. Naïve mice were age-matched, non-vaccinated controls. **(a)** Total p79-tetramer+ CD8 T cells per spleen of individual mice at d28 post-boost. Percentage of p79-dextramer+ CD8 T cells with markers of **(b)** short-lived effector cell (SLEC, KLRG1+CD127-) and **(c)** memory precursor effector cell subsets (MPEC, KLRG1-CD127+). **(d)** MPECs were further delineated into CD62L- effector and CD62L+ central MPECs for p79- dextramer+ CD8 T cells. **(e)** Percentage of Vβ4+ CD8 T cells in pooled spleens of mice with RDV-50.stop or RDV-50.stopΔM-M4 vaccination or infection with WT MHV68. For **(a-d)**, symbols represent individual mice, (N=5), and for **(e)** symbols represent pooled spleens from 5 mice.; bars and whiskers are mean +/- SD. *, p<0.05; **, p<0.01; ***, p<0.001 in Sidak's multiple comparisons test of one-way ANOVA **(a)**; in two-tailed unpaired t test **(b-d)** between the indicated groups.

**Supplementary Figure 3 Vaccination with RDV-50.stopΔM1-M4 reduces acute replication, latency and reactivation in C57BL/6 mice.** C57BL/6 mice either sham-vaccinated or prime-boost vaccinated IP with $1x10^6$ PFU of RDV-50.stopΔM1-M4 or RDV-50.stop followed by IP challenge with $5x10^3$ PFU of WT MHV68 at d28 post-boost. **(a)** Acute replication at d7 post-challenge determined by measuring infectious particles per ml lung homogenate. (N=5 mice per experiment); bars and whiskers represent mean +/- SD. ****, p<0.0001 in Sidak's multiple comparisons test of one-way ANOVA between the indicated groups. **(b)** The frequency of latency determined by limiting dilution nested

PCR of intact splenocytes for the viral genome at day d28 post-boost. **(c)** The frequency of explant reactivation determined by limiting dilution coculture of intact viable splenocytes on a monolayer of primary MEFs d28 post-boost. Disrupted splenocytes plated in parallel did not detect preformed infectious virus in the vaccinated animals. For **(b-c),** symbols represent pool of five individual mice per experiment.

**Supplementary Figure 4 Evaluation of T cell response to MHV68 in the spleens of vaccinated mice at seventeen days post-challenge with WT virus.** C57BL/6 mice were either sham-vaccinated or prime-boost vaccinated IP with $1x10^6$ PFU RDV-50.stopΔM1-M4 or RDV-50.stop followed by IN challenge with $1x10^3$ PFU WT MHV68 at d28 post-boost and analyzed d17 post-challenge. **(a)** Total p79-dextramer+ CD8 T cells per spleen of individual mice with the indicated vaccination regimen. Percentage of p79-dextramer+ CD8 T cells with markers of **(b)** short-lived effector cell (SLEC, $KLRG1^+CD127^-$) and **(c)** memory precursor effector cell subsets (MPEC, $KLRG1^-CD127^+$). **(d)** MPECs were further delineated into $CD62L^-$ effector and $CD62L^+$ central MPECs for p79-dextramer+ CD8 T cells. For each graph, symbols represent individual mice, (N=3-5); bars and whiskers are mean +/- SD. **, $p<0.01$ in Sidak's multiple comparisons test of one-way ANOVA between the indicated groups.

Supplemental Figures

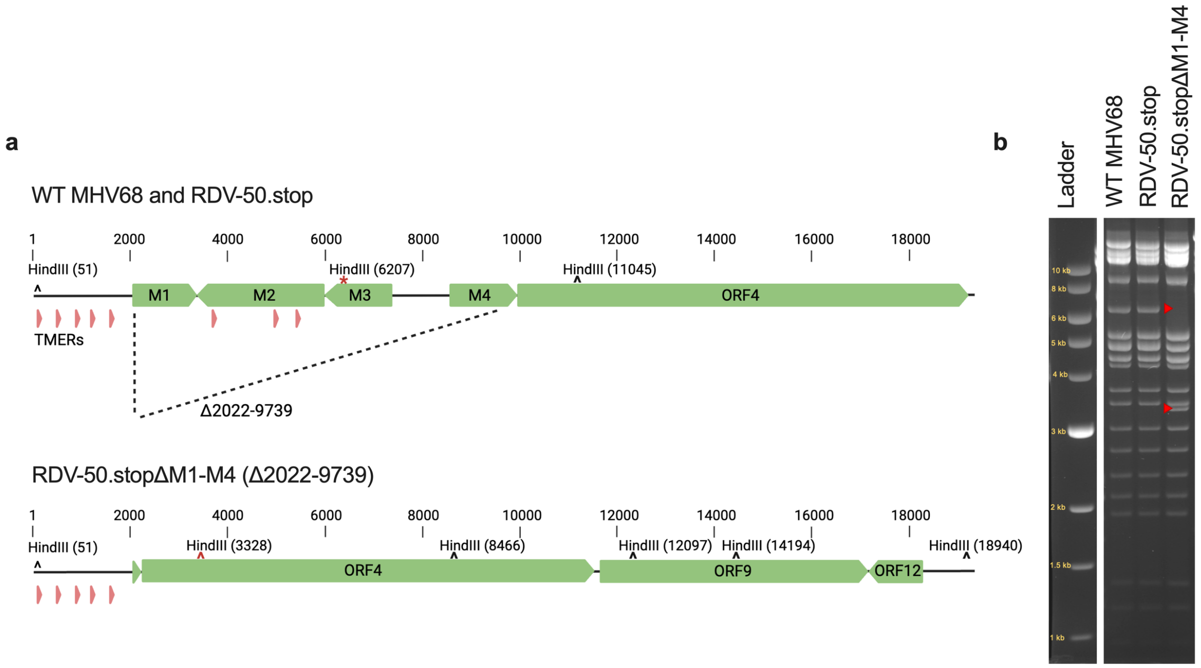


**Supplementary Figure 1 Generation of recombinant RDV-50.stopΔM1-M4 MHV68 upon deletion of unique M gene locus. (a)** Schematic describing the left end of WT MHV68 and the parental RDV-50.stop. M1- M4 genes and TMERs 6-8 (nucleotides 2022 to 9739) were deleted from RDV-50.stop to generate MHV68 RDV-50.stopΔM1-M4. '*' and '^' denote HindIII digestion sites. '*' indicates the site lost with the deletion of 2022 to 9739 bp. Figure made with BioRender. **(b)** In the HindIII restriction digest, WT MHV68 and RDV-50.stop has the expected 6156 bp fragment, while the RDV-50.stopΔM1-M4 genome has the expected loss of the 6156 bp fragment and gain of a 3277 bp fragment, indicated by red arrows.

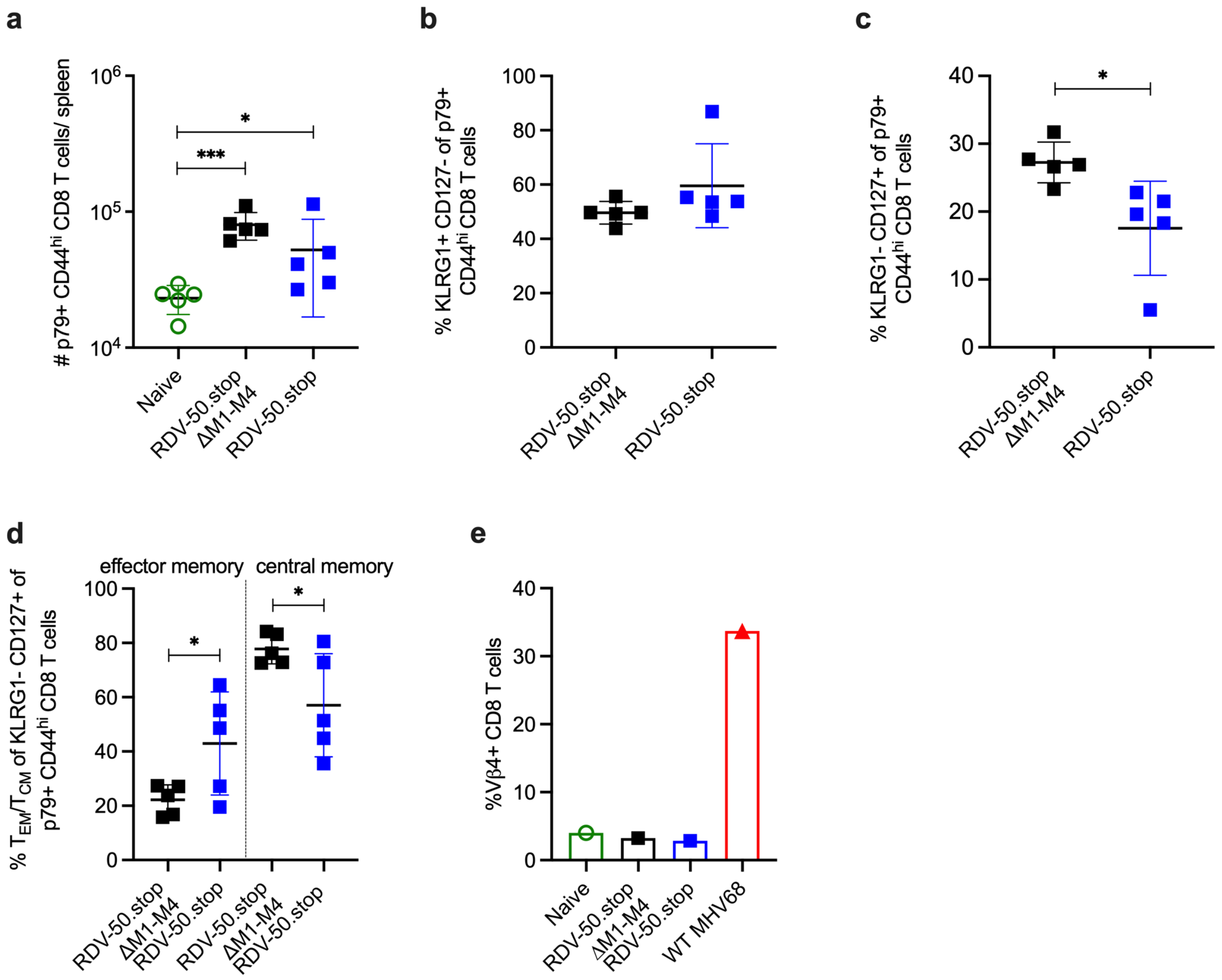


**Supplementary Figure 2 RDV-50.stopΔM1-M4 generates virus-specific CD8 T cell responses but does not induce M1 driven Vβ4+ CD8 T cells upon a prime-boost regimen in C57BL/6 mice.** C57BL/6 mice were either sham-vaccinated or prime-boost vaccinated IP with 1x10$^6$ PFU RDV-50.stopΔM1-M4 or RDV-50.stop. Naïve mice were age-matched, non-vaccinated controls. **(a)** Total p79-tetramer+ CD8 T cells per spleen of individual mice at d28 post-boost. Percentage of p79-dextramer+ CD8 T cells with markers of **(b)** short-lived effector cell (SLEC, KLRG1+CD127-) and **(c)** memory precursor effector cell subsets (MPEC, KLRG1-CD127+). **(d)** MPECs were further delineated into CD62L- effector and CD62L+ central MPECs for p79- dextramer+ CD8 T cells. **(e)** Percentage of Vβ4+ CD8 T cells in pooled spleens of mice with RDV-50.stop or RDV-50.stopΔM-M4 vaccination or infection with WT MHV68. For **(a-d)**, symbols represent individual mice, (N=5), and for **(e)** symbols represent pooled spleens from 5 mice.; bars and whiskers are mean +/- SD. *, p<0.05; **, p<0.01; ***, p<0.001 in Sidak's multiple comparisons test of one-way ANOVA **(a)**; in two-tailed unpaired t test **(b-d)** between the indicated groups.

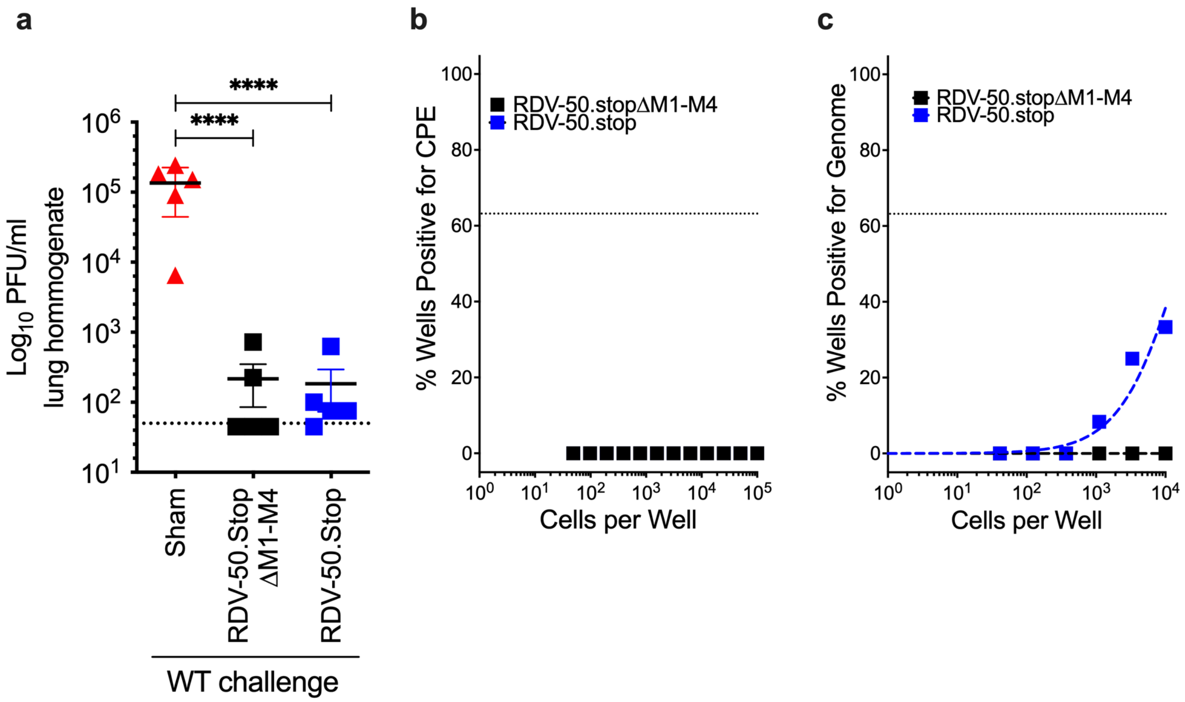


**Supplementary Figure 3 Vaccination with RDV-50.stopΔM1-M4 reduces acute replication, latency and reactivation in C57BL/6 mice.** C57BL/6 mice either sham-vaccinated or prime-boost vaccinated IP with $1x10^6$ PFU of RDV-50.stopΔM1-M4 or RDV-50.stop followed by IP challenge with $5x10^3$ PFU of WT MHV68 at d28 post-boost. **(a)** Acute replication at d7 post-challenge determined by measuring infectious particles per ml lung homogenate. (N=5 mice per experiment); bars and whiskers represent mean +/- SD. ****, $p<0.0001$ in Sidak's multiple comparisons test of one-way ANOVA between the indicated groups. **(b)** The frequency of latency determined by limiting dilution nested PCR of intact splenocytes for the viral genome at day d28 post-boost. **(c)** The frequency of explant reactivation determined by limiting dilution coculture of intact viable splenocytes on a monolayer of primary MEFs d28 post-boost. Disrupted splenocytes plated in parallel did not detect preformed infectious virus in the vaccinated animals. For **(b-c),** symbols represent pool of five individual mice per experiment.

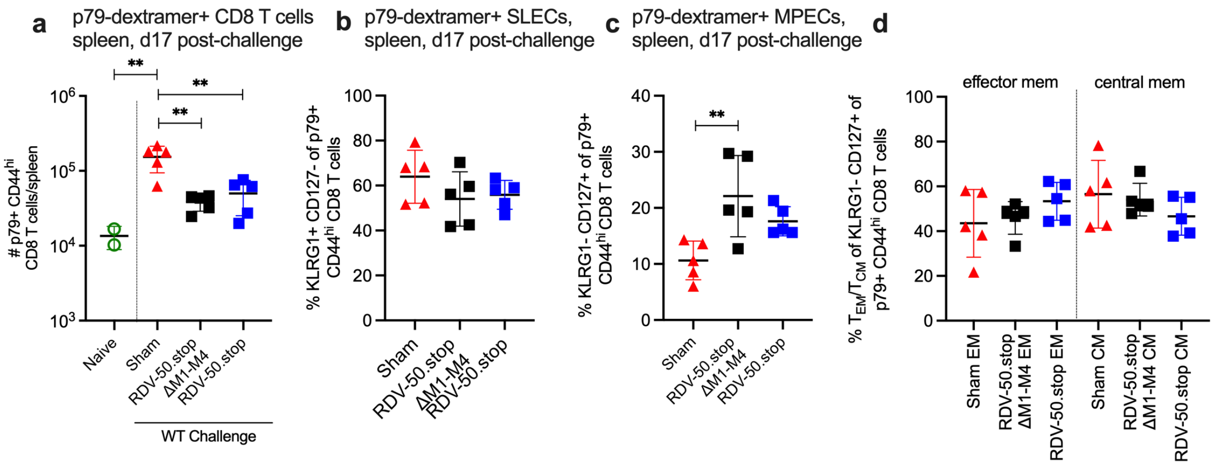


**Supplementary Figure 4 Evaluation of T cell response to MHV68 in the spleens of vaccinated mice at seventeen days post-challenge with WT virus.** C57BL/6 mice were either sham-vaccinated or prime-boost vaccinated IP with $1x10^6$ PFU RDV-50.stopΔM1-M4 or RDV-50.stop followed by IN challenge with $1x10^3$ PFU WT MHV68 at d28 post-boost and analyzed d17 post-challenge. **(a)** Total p79-dextramer+ CD8 T cells per spleen of individual mice with the indicated vaccination regimen. Percentage of p79-dextramer+ CD8 T cells with markers of **(b)** short-lived effector cell (SLEC, $KLRG1^+CD127^-$) and **(c)** memory precursor effector cell subsets (MPEC, $KLRG1^-CD127^+$). **(d)** MPECs were further delineated into $CD62L^-$ effector and $CD62L^+$ central MPECs for p79-dextramer+ CD8 T cells. For each graph, symbols represent individual mice, (N=3-5); bars and whiskers are mean +/- SD. **, p<0.01 in Sidak's multiple comparisons test of one-way ANOVA between the indicated groups.
